# Heat-inactivated modified vaccinia virus Ankara boosts Th1-biased cellular and humoral immune responses as a vaccine adjuvant by activating the STING-mediated cytosolic DNA-sensing pathway

**DOI:** 10.1101/2021.07.24.453449

**Authors:** Ning Yang, Aitor Garzia, Cindy Meyer, Thomas Tuschl, Taha Merghoub, Jedd D. Wolchok, Liang Deng

**Affiliations:** Dermatology Service, Department of Medicine, Memorial Sloan Kettering Cancer Center, New York, NY, 10065, USA; Laboratory of RNA Molecular Biology, The Rockefeller University, New York, NY, 10065, USA; Immuno-oncology service, Human Oncology and Pathogenesis Program; Memorial Sloan Kettering Cancer Center, New York, NY 10065, USA; Parker Institute for Cancer Immunotherapy, Memorial Sloan Kettering Cancer Center, New York, NY, USA; Weill Cornell Medical College, New York, NY, USA

**Keywords:** Vaccinia virus, poxvirus, vaccine adjuvant, dendritic cell maturation, STING (Stimulator of interferon genes), the cytosolic-DNA-sensing pathway, neoantigen, SARS-CoV2

## Abstract

**Background:** Protein or peptide-based subunit vaccines are promising platforms for combating human cancers and infectious diseases. However, one primary concern regarding subunit vaccines is the relatively weak immune responses induced by proteins or peptides. Therefore, developing novel and effective vaccine adjuvants is critical for the success of subunit vaccines. Modified vaccinia virus (MVA) is a safe and effective vaccine against smallpox and monkeypox. In this study, we explored the potential of heat-inactivated MVA (heat-iMVA) as a novel vaccine adjuvant.

**Methods:** We co-administered heat-iMVA with a model antigen, chicken ovalbumin (OVA), either intramuscularly or subcutaneously twice, two weeks apart, and analyzed anti-OVA specific CD8^+^ and CD4^+^ T cells in the spleens and skin draining lymph nodes (dLNs) and serum anti-OVA IgG1 and IgG2c antibodies. We also compared the adjuvanticity of heat-iMVA with several known vaccine adjuvants, including complete Freund’s adjuvant (CFA) and AddaVax, an MF59-like preclinical grade nano-emulsion. In addition, we tested whether co-administration of heat-iMVA plus tumor neoantigen peptides or irradiated tumor cells improves antitumor efficacy in a B16-F10 therapeutic vaccination model. Using Stimulator of Interferon Genes (STING) or Batf3-deficient mice, we evaluated the contribution of the STING pathway and Batf3-dependent CD103^+^/CD8α DCs in heat-iMVA-induced immunity.

**Results:** Co-administration of protein- or peptide-based immunogens with heat-iMVA dramatically enhances Th1-biased cellular and humoral immune responses. This adjuvant effect of heat-iMVA is dependent on the STING-mediated cytosolic DNA-sensing pathway, and the antigen-specific CD8^+^ T cell response requires Batf3-dependent CD103^+^/CD8α^+^ dendritic cells (DCs). Heat-iMVA infection of bone marrow-derived DCs (BMDCs) promoted antigen cross-presentation, whereas live MVA infection did not. RNA-seq analyses revealed that heat-iMVA is a more potent activator of the STING pathway than live MVA. Additionally, combining tumor neoantigen peptides or irradiated tumor cells with heat-iMVA delayed tumor growth and extended the median survival in B16-F10 therapeutic vaccination models.

**Conclusions:** Heat-iMVA induces type I interferon (IFN) production and antigen cross-presentation via a STING-dependent mechanism in DCs. Co-administration of heat-iMVA with peptide antigen generates strong Th1-biased cellular and humoral immunity. Collectively, our results demonstrate that heat-iMVA is a safe and potent vaccine adjuvant.

## BACKGROUND

Discoveries of cancer neoantigens generated by somatic mutations in cancer cells have brought excitement and renewed interest in cancer vaccines ^1–5^ and personalized neoantigen peptide vaccination has shown promising results in clinical trials ^2 6 7^. However, recombinant protein or peptide vaccines usually generate weak immune responses, safe and effective vaccine adjuvants that boost vaccine efficacy are urgently needed.

Licensed vaccine adjuvants include inorganic aluminum salts (alum), the oil-in-water emulsion MF59, monophosphoryl lipid A (MPL) absorbed on aluminum salts (AS04), and the toll-like receptor 9 (TLR9) agonist CpG 1018 ^8^. In addition to TLR agonists, agents that activate the cytosolic pattern recognition receptors, for example, stimulator of interferon genes (STING) agonists, have also been explored as vaccine adjuvants ^9 10^. It has been postulated that vaccine adjuvants that mimic natural infection might elicit potent and durable immune responses via the activation of innate immune-sensing pathways ^11^.

Vaccinia virus (VACV) belongs to the poxvirus family, and modified vaccinia virus Ankara (MVA) is a highly attenuated vaccinia strain, a safe and effective second-generation smallpox vaccine and a vaccine vector against other infectious agents ^12–19^. We have previously shown that wild-type vaccinia (WT VACV) infection of bone marrow-derived dendritic cells (BMDCs) fails to induce type I IFN production. By contrast, MVA infection induces IFN production via the cGAS/STING-mediated cytosolic DNA-sensing pathway ^20^.

Vaccinia virus encodes many immunomodulatory genes to evade the host immune system ^21–23^. Inactivation of WT VACV or MVA, by heating MVA at 55°C for 1h, reduces infectivity by 1000-fold and much more potently induces type I IFN production than live viruses in conventional DCs (cDCs) or plasmacytoid DCs (pDCs) ^24–26^. Based on its safety and immune-stimulating features, we hypothesized that heat-inactivated vaccinia or MVA could act as vaccine adjuvant. Here, we show that the heat-inactivated MVA (heat-iMVA) can boost the T and B cell responses of subunit vaccines. Furthermore, co-administration of heat-iMVA with tumor neoantigen peptides delays tumor growth and prolongs mouse survival in a syngeneic B16-F10 melanoma model. In summary, our results provide proof-of-concept for heat-iMVA as a vaccine adjuvant against infectious diseases and cancers.

## MATERIALS AND METHODS

### Study design

We used intramuscular and subcutaneous vaccinations to compare the adjuvanticity of heat-iMVA with known vaccine adjuvants, CFA and Addavax. Controls groups with PBS mock vaccination were used. OVA was used as a model antigen. In most of the experiments, female C57BL/6J mice were used. The sample size calculation was based on expected immune adjuvant effects of known adjuvants and preliminary results with heat-iMVA as a vaccine adjuvant, variability of the measurements, and a target power of 95%. Randomization was performed to minimize confounders among the treatment and control groups. The researcher who performed the outcome assessment and the data analysis were not aware of the group allocation. Outcome measures include tumor volumes and mice survival.

### Mice

Female C57BL/6J mice between 6 and 8 weeks of age were purchased from the Jackson Laboratory and were used for vaccination experiments and for the preparation of bone marrow-derived dendritic cells (BMDCs). Batf3^-/-^, STING^Gt/Gt^, and cGAS^-/-^ mice were generated in the laboratories of Kenneth Murphy, Russell Vance, and Herbert (Skip) Virgin, respectively. Mice deficient for IFNα/*β* receptor (IFNAR^-/-^) were provided by Eric Pamer. IFNAR^-/-^OT-1 mice were generated by crossing IFNAR^-/-^ mice and OT-1 transgenic mice for several generations. All mice were maintained in the animal facility at the Sloan Kettering Cancer Institute. All procedures were performed in strict accordance with the recommendations in the *Guide for the Care and Use of Laboratory Animals* of the National Institute of Health. The protocol was approved by the Committee on the Ethics of Animal Experiments of Sloan-Kettering Cancer Institute. We used the ARRIVE reporting guidelines ^27^

### Cell lines and primary Cells

BHK-21 (baby hamster kidney cell, ATCC CCL-10) cells were used to propagate the MVA virus. The procedure for the generation of GM-CSF-BMDCs and Flt3L-BMDCs have been described ^26^. HEK293T cell line expressing human ACE2 (hACE2) were generated by transduction with vesicular stomatitis virus (VSV) G protein-pseudotyped murine leukemia viruses (MLV) containing pQCXIP-hACE2-c9 as described ^28^. The murine melanoma cell line B16-F10 was originally obtained from I. Fidler (MD Anderson Cancer Center). B16-GM-CSF cells were generated by Glenn Dranoff ^29^.

### Viruses

MVA virus was kindly provided by Gerd Sutter (University of Munich). Heat-iMVA was generated by incubating purified MVA virus at 55 °C for 1 hour ^25^. SARS-CoV-2 pseudoviruses were produced using a method as described previously ^30^. Briefly, HEK293T cells were co-transfected with pQCXIG-SARS-CoV-2-Spike, pMD2.G (VSV-G) and a gag/pol expression plasmid. At 48 h post-transfection, virus supernatants were harvested and filtered through a 0.45- μm filter and stored at −80 °C.

### Reagents

EndoFit Ovalbumin, CFA, AddaVax, and poly(I:C) were purchased from InvivoGen. The SARS-CoV-2 spike protein was purchased from RayBiotech. Alexa FluorTM 647 conjugated OVA was purchased from Thermo Fisher. B16-F10 tumor neoantigen peptides were synthesized by GenScript (Piscataway, NJ). The sequences are as follows: M27: REGVELCPGNKYEMRRHGTTHSLVIHD; M30: PSKPSFQEFVDWENVSPELNSTDQPFL; M48: SHCHWNDLAVIPAGVVHNWDFEPRKVS using the mutation information described ^4^.

### OVA vaccination procedure

WT C57BL/6J mice were anesthetized and vaccinated initially on day 0 and boosted on day 14 with either OVA (10 µg) alone, or OVA (10 µg) + heat-iMVA (an equivalent of 10^7^ pfu) in a volume 100 µl intramuscularly (IM) or subcutaneously (SC). Mice were euthanized on day 21. Spleens, draining lymph nodes (dLNs), and blood were collected for analyzing OVA-specific CD8^+^, CD4^+^, and B cell responses. In some cases, OVA proteins were mixed with CFA or AddaVax. In some cases, STING^Gt/Gt,^ Batf3^-/-^, and age-matched WT C57BL/6J mice were vaccinated with OVA + heat-iMVA as described above.

### SARS-CoV-2 spike protein vaccination procedure

4-5 mice in each group were anesthetized and vaccinated with SARS-CoV-2 recombinant spike protein (1 µg) alone or with spike (1 μg) + heat-iMVA in a volume 100 µl intramuscularly on day 0 and boosted on day 21. Mice were euthanized on day 28. Spike-specific immunoglobulin G1 (IgG1) or immunoglobulin G2c (IgG2c) titers in the serum from PBS, spike alone, or spike + heat-iMVA-vaccinated mice were determined by ELISA.

### Therapeutic vaccination using neoantigen peptides with or without adjuvants

B16-F10 melanoma cells (5 x 10^4^) were implanted intradermally into the shaved skin on the right flank of WT C57BL/6J mice. On day 3, 6, and 9, 4 groups of mice (10 mice in each group) were subcutaneously vaccinated at the left flanks with B16-F10 neoantigen peptide mix (M27, M30 and M48) (100 µg each) ^4^, with or without heat-iMVA or poly(I:C) (50 µg), or with PBS mock control. Mice were monitored daily, and tumor sizes were measured twice a week. Tumor volumes were calculated according to the following formula: *l* (length) x *w* (width) x *h* (height)/2. Mice were euthanized for signs of distress or when the diameter of the tumor reached 10 mm.

### Therapeutic vaccination using irradiated B16-GM-CSF whole-cell with and without adjuvants

B16-F10 melanoma cells (5 x 10^4^) were implanted intradermally into the shaved skin on the right flank of WT C57BL/6J mice. On day 3, 6, and 9, four groups of mice (10 mice in each group) were subcutaneously vaccinated at the left flanks with irradiated B16-GM-CSF (1 x 10^6^ cells after 150 Gy *γ*-irradiation) with or without heat-iMVA or poly(I:C), or with PBS mock control. Mice were monitored daily, and tumor sizes were measured twice a week.

### Flow cytometry analysis of antigen-specific T cells in the spleens and dLNs

Spleens and dLNs from vaccinated mice was collected and processed using Miltenyi GentleMACS™ Dissociator. Red blood cells were lysed using ACK buffer. For intracellular cytokine staining, splenic or lymph node single-cell suspensions were stimulated with 10 μg/ml peptides (OVA_257-264_ or OVA_323-339_). After 1 h of stimulation, GolgiPlug (BD Biosciences) (1:1000 dilution) was added and incubated for 12 h. Cells were then treated with BD Cytofix/Cytoperm™ kit prior to staining with respective antibodies for flow cytometry analyses. The antibodies used for this assay are as follows: BioLegend: CD3e (145-2C11), CD4 (GK1.5), CD8 (53-5.8), IFN-*γ* (XMG1.2).

### Antibodies titer determination by ELISA

ELISA was used to determine anti-OVA or anti-SARS-CoV-2 spike IgG titers. Briefly, 96-well microtiter plates (Thermo Fisher) were coated with 2.0 µg/mL of OVA (Invivogen) or SARS-CoV-2 spike protein (RayBiotech) overnight at 4°C. Plates were washed with 0.05% Tween-20 in PBS (PBST) and blocked with 1% BSA/PBS-T. Mouse serum samples were two-fold serially diluted in PBST, added to the blocked plates, and incubated at 37°C for 1 h. Following incubation, plates were washed with PBS-T and incubated with horseradish peroxidase (HRP)-conjugated goat anti-mouse IgG1 or goat anti-mouse IgG2c (Southern Biotech) for 1 h. Plates were washed with PBS-T and TMB substrate (BD Bioscience) was added. Reactions were stopped with 50 µl 2N H_2_SO4. Plates were read at OD 450 nm with a SpectraMax Plus plate reader (Molecular Devices). The antibody titer is defined as the dilution in which absorbance is more than 2.1 times of the blank wells.

### Flow cytometry analysis of migratory and skin LN-resident DCs after fluorescent-labeled OVA-647 vaccination with or without heat-iMVA

C57BL/6J mice were vaccinated intradermally with either Alexa Fluor 647-labeled OVA (OVA-647) alone or OVA-647 + heat-iMVA. Skin dLNs were harvested at 24 h post injection, digested with collagenase D (2.5 mg/ml) and DNase (50 µg/ml) at 37°C for 25 min before filtering through 70-µm cell strainer, and analyzed by flow cytometry for OVA-647 intensities and CD86 expression of the migratory DC and resident DC populations in the skin dLNs. The antigens and clone designations for the antibodies are as follows: BioLegend: CD11c (N418). CD11b (M1/70), MHC-II (M5/114.15.2), CD3e (145-2C11), CD8a (53-6.7); BD Biosciences: Siglec F (E50-2440), CD19 (1D3), CD49b (DX5), CD207 (81E2), Thermo Fisher: CD16/CD32 (93), CD103 (2E7), TER-119 (TER-119). Cells were analyzed on the BD LSR Fortessa or LSR II flow cytometer and data were analyzed with FlowJo software (version 10.5.3).

### RNA-seq analyses of GM-CSF-cultured BMDCs infected with live MVA vs. Heat-iMVA

GM-CSF-cultured BMDCs (1 x 10^6^) from WT or STING^Gt/Gt^ mice were infected with live MVA or heat-iMVA at a multiplicity of infection (MOI) of 10. Cells were collected at 2, 4, and 6 h post-infection. Total RNA was extracted using TRIzol (Thermo Fisher). Agilent 2100 Bioanalyzer at the Rockefeller University Genomics Resource Center was used to assess total RNA integrity and quantity. Samples with the RNA integrity number (RIN) > 9.5 were used. Oligo(dT)-selected RNA was converted into cDNA for RNA sequencing using the Illumina TruSeq RNA Sample Preparation Kit v2 according to the instructions of the manufacturer and sequenced on an Illumina HiSeq 2500 platform using 100 nt single-end sequencing. Reads were aligned against the mouse genome (Gencode, GRCm38) plus vaccinia genome (VACV-MVA) using TopHat v2.0.14 (http://tophat.cbcb.umd.edu/). The steps described followed the protocol by Trapnell and colleagues ^31^. Cufflinks v2.1.1 (http://cole-trapnell-lab.github.io/cufflinks/) was used for estimation of transcript abundance and differential expression analysis. Unsupervised hierarchical clustering was performed using Euclidean distance and complete linkage for columns (samples) and rows (mRNAs). For clarity, the row dendrograms were removed from the figures. The R packages pheatmap was used for data representation. Gene set enrichment analysis (GSEA) was conducted using the package fgsea with 1000 permutations, with reactome and MSigDB C7 signature sets.

### Antigen cross-presentation assay

GM-CSF-cultured or Flt3L-cultured BMDCs were infected or mock-infected with heat-iMVA at a MOI of 1 and then added OVA at indicated concentrations and incubated for 3 h. Cells were washed with fresh medium and co-cultured with carboxyfluorescein diacetate succinimidyl ester (CFSE)-labeled OT-1 for 3 days (BMDC:OT-1 T-cells =1:5). Flow cytometry was applied to measure CFSE intensities of OT-Ι cells.

WT or STING-deficient GM-CSF-cultured BMDCs were incubated with OVA in the presence or absence of either live MVA or heat-iMVA for 3 h. Cells were washed with fresh medium and co-cultured with OT-1 cells (BMDC to OT-1 T-cells ratio of 1:3) for 3 days. IFN-*γ* levels in the supernatants were determined by ELISA (R&D). OT-1 cells were purified from OT-1 transgenic mice using negative selection with CD8a^+^ T Cell Isolation Kit according to the manufacturer’s instructions (Miltenyi Biotec).

### SARS-CoV-2 pseudovirus neutralization assay

Serially diluted serum was pre-incubated with SARS-CoV-2 pseudovirus at room temperature (RT) for 30 mins, and the mixtures were added to 293T-hACE2. Medium was changed 2 h later. After 48 h, cells were fixed in 4% paraformaldehyde in PBS for 15 min at RT. Cell nuclei were stained with Hoechst 33258 (Sigma) in PBS for 10 min at RT. Images were captured using Zeiss Axio Observer 7 (Carl Zeiss) and analyzed with ZEN Imaging software (Carl Zeiss) and Image J (Fiji). GFP expression in pseudovirus-infected cells were determined using the BD LSR Fortessa flow cytometer and data were analyzed using FlowJo software (version 10.5.3).

### Statistics

Two-tailed, unpaired, Student’s *t* test was used for comparisons of two independent groups in the studies. Survival data were analyzed by log-rank (Mantel-Cox) test. The *P* values deemed significant are indicated in the figures as follows: \**P* < 0.05; \*\**P* < 0.01; \*\*\**P* < 0.001; \*\*\*\**P* < 0.0001. Statistical analyses were performed on the Prism GraphPad Software. The numbers of animals included in the study are discussed in each figure legend.

## RESULTS

### Co-administration of chicken ovalbumin (OVA) with heat-iMVA enhances the generation of OVA-specific cellular and humoral immune responses in mice

We prime-immunized mice intramuscularly (IM) with the model antigen OVA with or without heat-iMVA, followed by a boost-immunization two weeks later, and euthanized them one week after the boost vaccination. IM co-administration of heat-iMVA with OVA increased splenic anti-OVA IFN-γ ^+^CD8^+^ and CD4^+^ T-cells compared with OVA alone (figure 1A-D). We also observed a stronger induction of anti-OVA IFN-γ ^+^CD8^+^ and CD4^+^ T-cells compared with OVA alone in the skin draining lymph nodes (dLNs) (Supplemental figure. 1A-D). In addition, the combination of heat-iMVA plus OVA induced stronger IgG1 production than OVA alone (figure 1E). IgG2c antibody titers were upregulated by 25-fold in the OVA plus heat-iMVA group than OVA alone (figure 1F), suggesting that prime-boost vaccination with the combined heat-iMVA/OVA induced stronger Th1 immune responses.

**Figure 1.**
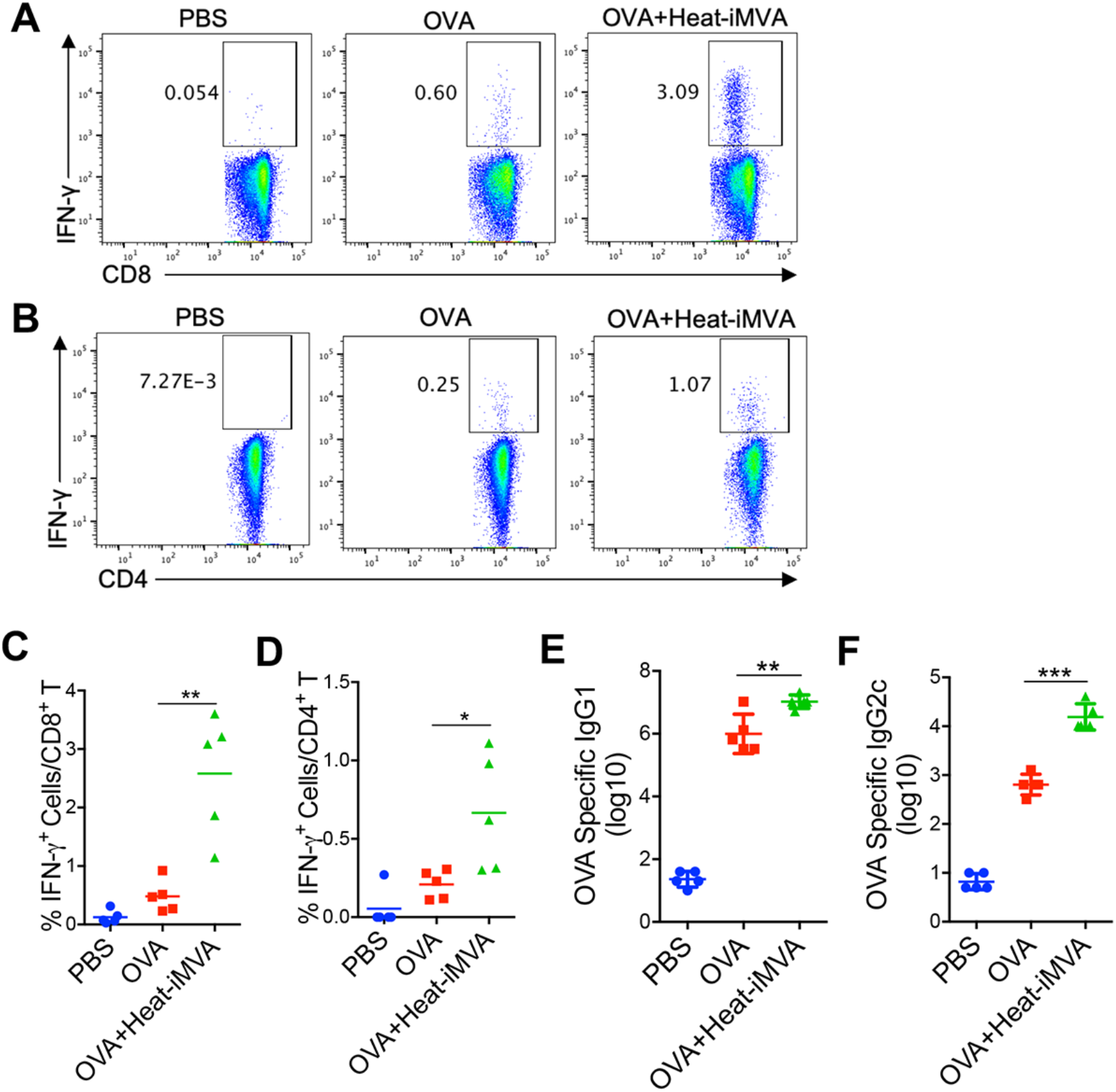
Co-administration of heat-inactivated MVA (heat-iMVA) enhances antigen-specific T cell and antibody responses after intramuscular (IM) vaccination with chicken ovalbumin (OVA). WT C57BL/6J mice were vaccinated on day 0 and day 14 with OVA (10 μg) or OVA (10 μg) plus heat-iMVA (an equivalent amount of 10^7^ pfu/mouse) intramuscularly. On day 21, splenocytes (A, B, C, D) were stimulated with OVA_257-264_ (A, C) or OVA_323-339_ peptide (B, D). The expression of IFN-γ by CD8^+^ T cells or CD4^+^ T was measured by flow cytometry. (E-F) OVA-specific immunoglobulin G1 (IgG1) or OVA-specific immunoglobulin G2c (IgG2c) titers in the serum from PBS, OVA, or OVA + heat-iMVA-vaccinated mice were determined by ELISA. Data are represented as mean ± SEM (*n* = 3-5; \**P* < 0.05, \*\**P* < 0.01 and \*\*\**P* < 0.001; unpaired *t* test). Data are representative of three independent experiments.

### Heat-iMVA promotes more robust Th1 responses and IgG2c production compared with complete Freund adjuvant (CFA) and AddaVax

Next, we compared the adjuvanticity of heat-iMVA with other well-known vaccine adjuvants. CFA comprises heat-killed *Mycobacterium tuberculosis* in non-metabolizable oils and also contains ligands for TLR2, 4, and 9. Injection of antigen with CFA induces a Th1-dominant immune response ^32^. Although CFA’s use in humans is currently impermissible due to its toxicity profile, it is commonly used in animal studies because of its strong adjuvant effects. Subcutaneous co-administration of OVA with heat-iMVA induced higher levels of antigen-specific CD8^+^ and CD4^+^ T-cells than OVA plus CFA in the spleens of vaccinated mice (figure 2A and 2C). Similarly, co-administration of OVA with heat-iMVA induced stronger OVA-specific CD8^+^ and CD4^+^ T cell responses in skin dLNs compared with OVA + CFA (Supplemental figure. 2A and 2B). Serum IgG1 titers from OVA + CFA-immunized mice were 6-fold higher than those in the serum from OVA + heat-iMVA-immunized mice (Fig 2C), whereas serum IgG2c titers from OVA + CFA-immunized mice were 10-fold lower than those in the serum of OVA + heat-iMVA-immunized mice (Fig 2D). These results indicate that co-administration of OVA plus Heat-iMVA promotes stronger Th1-biased humoral immunity than OVA plus CFA.

**Figure 2.**
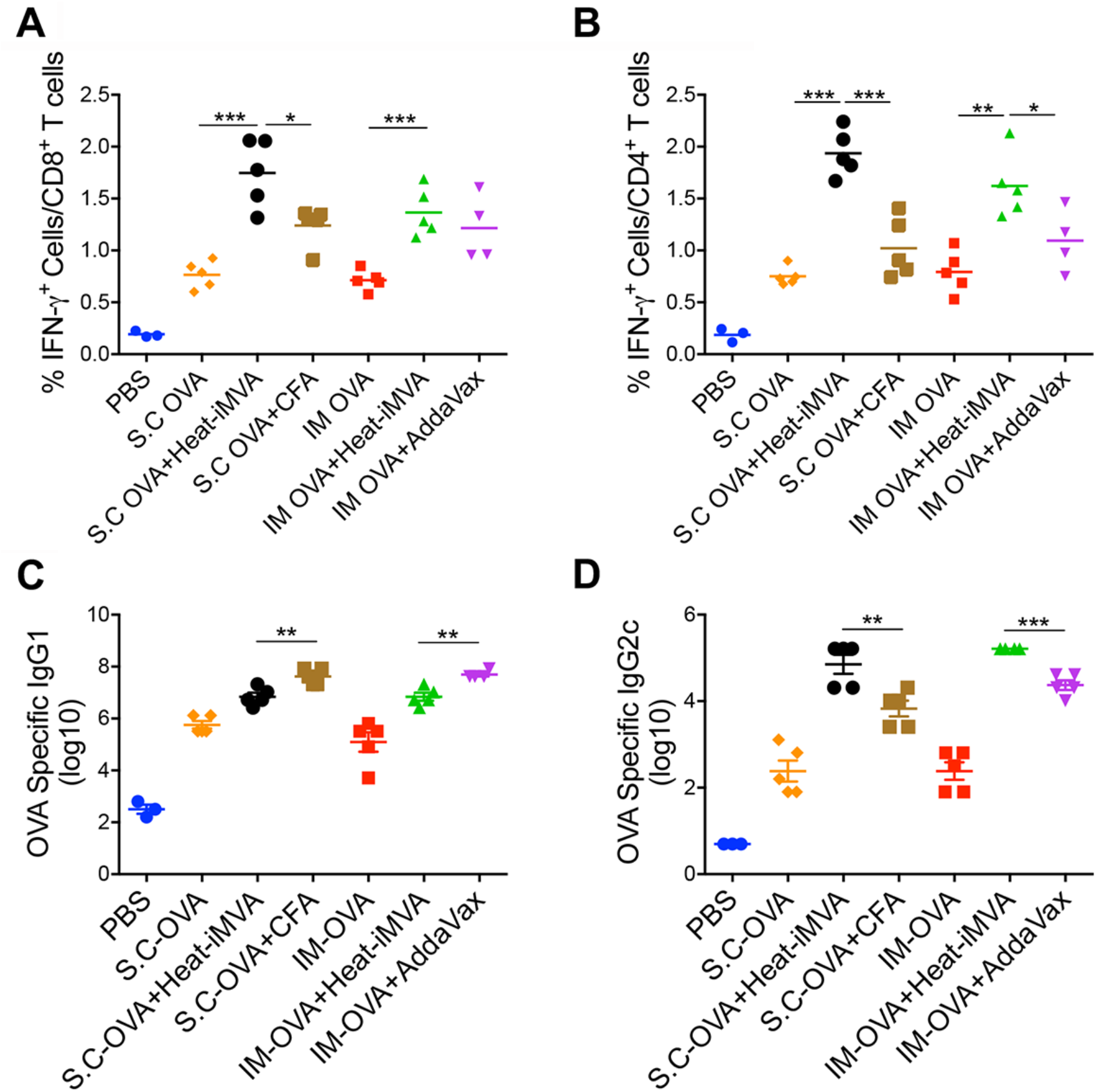
Heat-iMVA promotes stronger antigen-specific Th1 responses and IgG2c production compared with complete Freund adjuvant (CFA) and AddaVax after cutaneous vaccination. Antigen-specific T cell and antibodies responses were measured after intramuscular (IM) or subcutaneous (SC) vaccination on day 0 and day 14 with OVA (10 μg) in the presence or absence of heat-iMVA (an equivalent amount of 10^7^ pfu) in C57BL/6J mice. (A-B) On day 21, splenocytes were stimulated with OVA_257-264_ or OVA_323-339_. The expression of IFN-γ by CD8^+^ or CD4^+^ T T cells was measured by flow cytometry. (C-D) On day 21, OVA-specific immunoglobulin G1 (IgG1) or OVA-specific immunoglobulin G2c (IgG2c) titers in the were determined by ELISA. Data are represented as mean ± SEM (*n* = 3-5; \**P* < 0.05, \*\**P* < 0.01 and \*\*\**P* < 0.001; unpaired *t* test). Data are representative of two independent experiments.

MF59, a squalene-based oil-in-water vaccine adjuvant in the inactivated influenza vaccine Fluad, is also the adjuvant in subunit vaccines against SARS-CoV-2 ^33^. AddaVax is an MF59-like preclinical grade nano-emulsion that induces both Th1-cellular immune responses and Th2-biased humoral responses ^34^. Intramuscular (IM) vaccination with OVA plus heat-iMVA induced CD8^+^ T cell responses similar to OVA plus AddaVax, however, the former combination promoted higher CD4^+^ T cell responses than the latter (figure 2A and 2B). IM vaccination of OVA plus heat-iMVA induced 7-fold higher OVA-specific IgG2c titers (figure 2C) and 7-fold lower OVA-specific IgG1 than OVA plus AddaVax (figure 2D), suggesting that co-administration of the antigen plus heat-iMVA more potently induces antigen-specific Th1-biased cellular and humoral immune responses compared with combining the antigen with AddaVax. Overall, SC or IM co-administration of OVA with heat-iMVA generated similar cellular and humoral immune responses to OVA (figure 2A-D; Supplemental figure. 2A and 2B).

### The role of CD103^+^ /CD8α^+^ DCs and the STING pathway on heat-iMVA-induced vaccine adjuvant effects

BATF3 is a transcription factor critical for the development of CD103^+^/CD8α^+^ lineage DCs, which plays an essential role in cross-presenting viral and tumor antigens ^35^. Our results showed that the percentage of anti-OVA IFN-*γ*^+^ T-cells among splenic CD8^+^ T-cells induced by heat-iMVA was reduced in Batf3^-/-^ mice (figure 3A), whereas the generation of splenic anti-OVA IFN-*γ*^+^ CD4^+^ T-cells seemed unaffected (figure 3B), with minimal effects on the IgG1 and IgG2c production (figure 3C and 3D). These results support a role for Batf3-dependent CD103^+^/CD8α^+^ DCs in cross-presenting OVA antigen to generate OVA-specific splenic CD8^+^ T-cells in our vaccination model. STING agonist cGAMP can be used as a vaccine adjuvant ^36^. Here, we observed that the percentage of splenic anti-OVA IFN-*γ*^+^ CD8^+^ and CD4^+^ T-cells induced by heat-iMVA decreased in STING^Gt/Gt^ mice (figure 3A and 3B). Moreover, serum IgG2c titers were reduced by 10-fold in the STING^Gt/Gt^ mice vaccinated with OVA + heat-iMVA compared with immunized WT mice (figure 3D), while serum IgG1 titers did not significantly differ between the two groups (figure 3C). These results demonstrate that the cGAS/STING-mediated cytosolic DNA-sensing pathway plays a critical role in the vaccine adjuvant effects of heat-iMVA.

**Figure 3.**
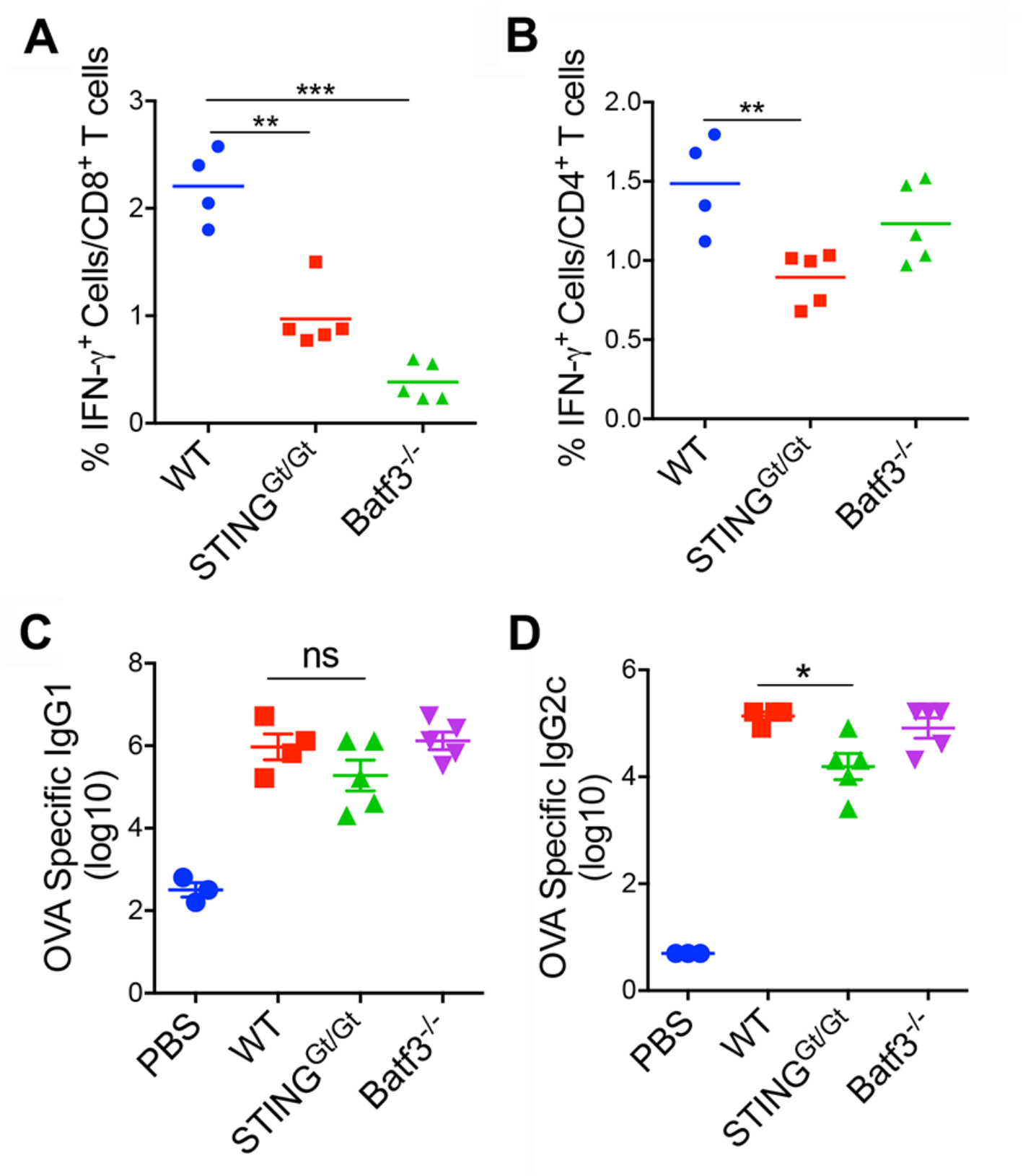
CD103^+^ DC and the cGAS/STING pathway contribute to heat-iMVA adjuvanticity. STING^Gt/Gt^, Batf3^-/-^, or age-matched WT C57BL/6J mice were intramuscularly vaccinated on day 0 and day 14 with OVA (10 µg) + heat-iMVA (an equivalent of 10^7^ pfu). (A-B) On day 21, mice were euthanized, and spleens and blood were collected. splenocytes were stimulated with OVA_257-264_ or OVA_323-339_. The expression of IFN-γ by CD8^+^ or CD4^+^ T cells was measured by flow cytometry. (C-D) On day 21, OVA-specific immunoglobulin G1 (IgG1) or OVA-specific immunoglobulin G2c (IgG2c) titers in the serum were determined by ELISA. Data are represented as mean ± SEM (*n* = 3-5; \**P* < 0.05, \*\**P* < 0.01 and \*\*\**P* < 0.001; unpaired *t* test). Data are representative of three independent experiments.

### Co-incubation of BMDCs with heat-iMVA and soluble OVA enhances antigen cross-presentation and proliferation of OT-I T-cells in vitro

Pre-incubation of GM-CSF-cultured BMDCs with heat-iMVA followed by pulsing with OVA enhanced the capacity of BMDCs to stimulate the proliferation of OT-I T-cells at all OVA concentrations, as indicated by CSFE dilution in the dividing cells (figure. 4A and 4B). Heat-iMVA potently stimulated Flt3L-BMDCs’ abilities to cross-present OVA and promote the proliferation of OT-I cells, even at an OVA concentration of 0.01 mg/ml (figure. 4C and 4D). In addition, GM-CSF-cultured BMDCs pre-infected with heat-iMVA were more potent in cross-presenting OVA antigen and stimulating IFN-*γ* secretion from proliferated and activated OT-I cells than BMDCs pre-infected with live MVA (figure 4E). IFN-*γ* levels were much lower in STING-deficient DCs than in WT DCs pretreated with heat-IMVA plus OVA and co-cultured with OT-I cells (figure 4E). Similar reduction of IFN-*γ* secretion by OT-1 cells were obtained in cGAS-deficient GM-CSF-cultured BMDCs compared with WT DCs (figure 4G). To test whether the STING/IFNAR pathway is important for heat-iMVA-induced antigen cross-presentation in CD103^+^ DCs, we sorted CD103^+^ DCs from Flt3L-cultured BMDCs from WT, STING^Gt/Gt^, or IFNAR^-/-^ mice, and preformed antigen cross-presentation assay. IFN-*γ* levels were much lower in heat-iMVA-infected STING-deficient or IFNAR^-/-^ CD103^+^ DCs than WT CD103^+^ DCs (figure 4F). These results indicate that the STING/IFNAR pathway plays a vital role in heat-iMVA-induced, CD103^+^ DCs-mediated antigen cross-presentation and antigen-specific T cell proliferation and activation. Finally, we tested whether IFNAR signaling on OT-I cells plays a role in T cell activation and we observed that IFNAR^-/-^ OT-1 cells secreted lower amount of IFN-*γ* when stimulated with OVA-pulsed heat-iMVA-treated GM-CSF-BMDCs (figure 4G).

**Figure 4.**
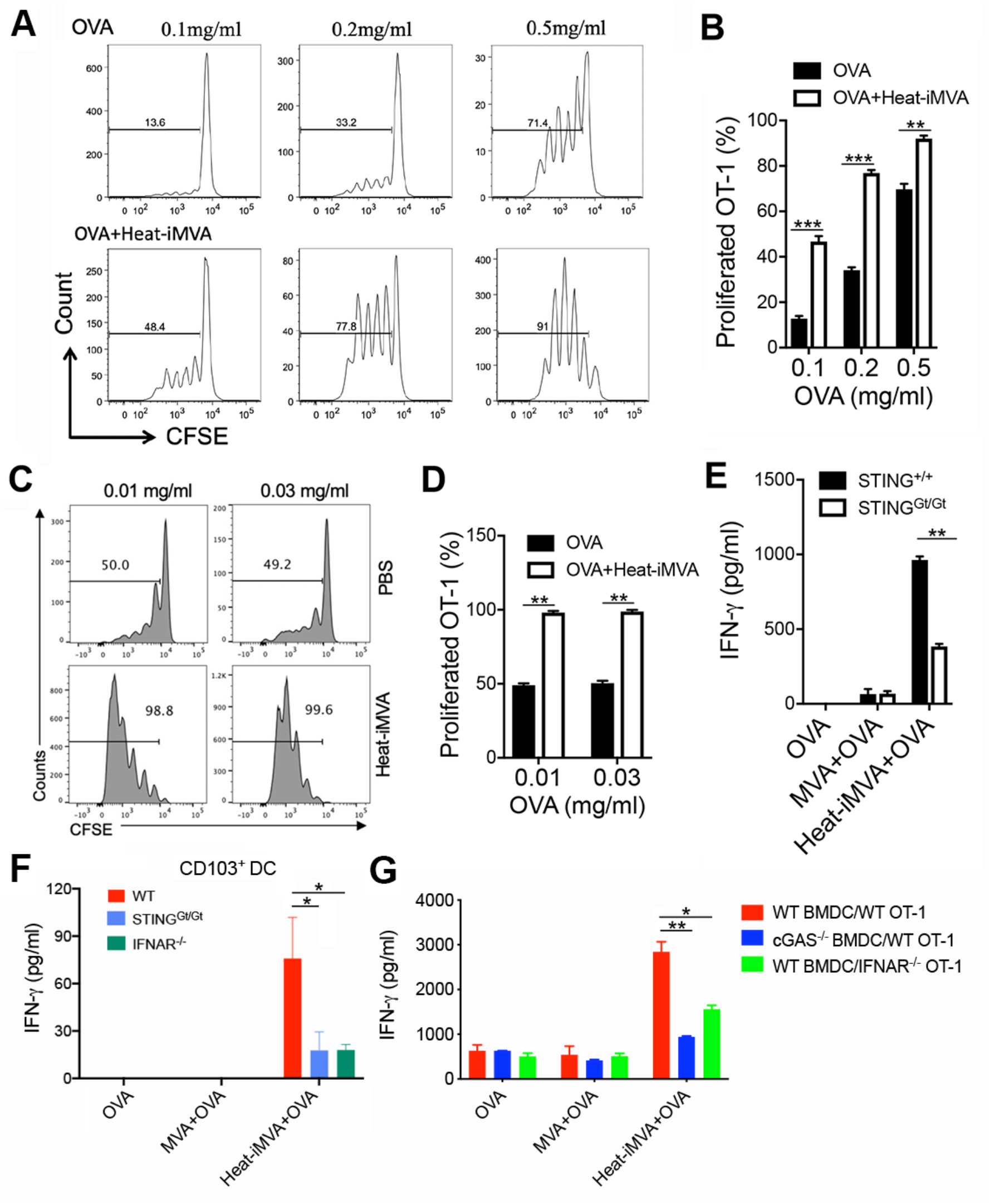
Heat-iMVA promotes OT-I cell activation and proliferation mediated by OVA cross-presentation by dendritic cells in vitro. (A, B, C, D) Proliferation of CFSE-labeled OT-Ι T cells after incubation with GM-CSF-cultured BMDCs (A, B) or FLT3L-cultured dendritic cells (C, D) pulsed with OVA in the presence or absence of heat-iMVA. BMDCs were incubated with or without heat-iMVA, and then co-cultured with CFSE-labeled OT-Ι cells for 3 days. (E) IFN-*γ* secretion from OT-Ι T cells after incubation with GM-CSF-cultured WT or STING^Gt/Gt^ BMDCs pulsed with OVA in the presence or absence of live MVA or heat-iMVA. (F) IFN-*γ* secretion from OT-Ι T cells after incubation with sorted CD103^+^ DCs from WT, STING^Gt/Gt^, or IFNAR^-/-^ Flt3L-cultured BMDCs pulsed with OVA in the presence or absence of live MVA or heat-iMVA. (G) IFN-*γ* secretion from WT or IFNAR^-/-^ OT-Ι T cells after incubation with OVA-pulsed GM-CSF-cultured DCs from WT or cGAS^-/-^ mice with or without live MVA or heat-iMVA. Data are represented as mean ± SEM (*n* = 3-5; \*\**P* < 0.01 and \*\*\**P* < 0.001; unpaired *t* test). Data are representative of three independent experiments.

### Heat-iMVA infection of BMDCs induces STING-dependent IFN and inflammatory cytokine responses

We performed RNA-seq analyses of BMDCs from WT and STING^Gt/Gt^ mice infected with either live MVA or heat-iMVA for 2, 4, and 6 h. Our results showed several patterns of gene expression induced by MVA and heat-iMVA: (i) Infection with live MVA, but not heat-iMVA, induced a subset of host genes in a STING-independent manner, thus indicating gene induction by a live virus infection (figure 5A, marked as a1-2); (ii) Heat-iMVA induced higher levels of a large subset of IFN-regulated genes than live MVA, which were mainly dependent on STING (figure 5A, marked as b1-3); and (iii) Heat-iMVA infection induced higher levels of a relatively small subset of genes than MVA, which were independent of STING (figure 5A, marked as c). Selected examples of genes in each category are shown in figure 5B. For example, heat-iMVA infection triggered higher levels of IFN-inducible genes than MVA, including *Ifih1 (MDA5), Ddx58 (RIG-1), Oasl2, Oas3, TLR3, Nod1, Ifna4, Ifnb1, Ccl5, Cxcl9, Cxcl10,* and members of the guanylate binding protein (Gbp) family, as largely dependent on STING (figure 5B, marked as b). These results indicate that the activation of the cytosolic DNA-sensing pathway by heat-iMVA also triggers the up-regulation of genes involved in the cytosolic RNA-sensing pathway in addition to other antiviral genes, thereby strengthening a broad range of innate immunity.

**Figure 5.**
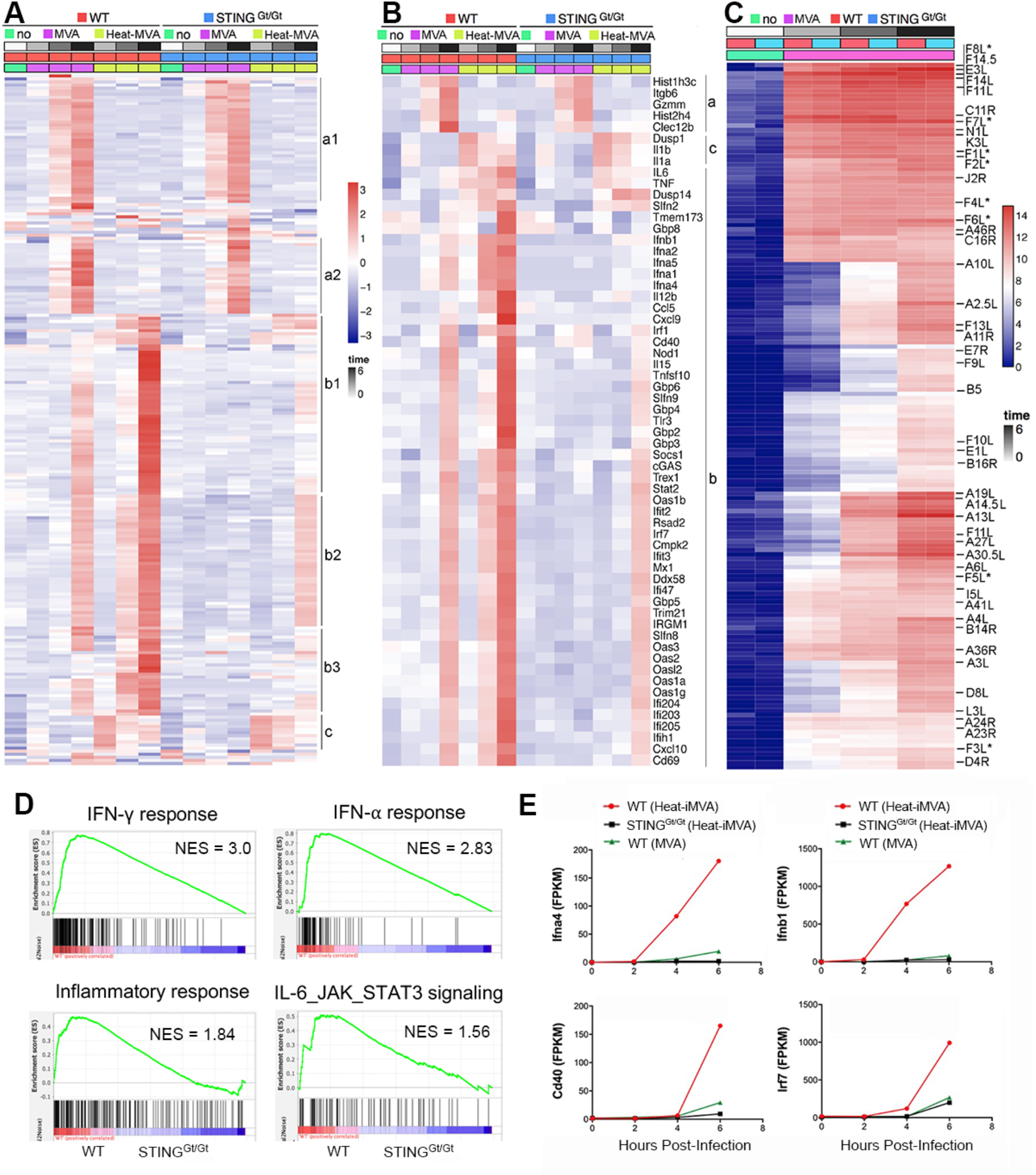
Time-resolved transcriptome profiling of WT or STING^Gt/Gt^ BMDCs infected with either live MVA or heat-iMVA. (A) A heat map of a one-way hierarchical clustering analysis of the top 200 genes ranked by Z-score of log2RPKM, indicating genes that exhibited the most statistically significant changes in gene expression over the course of the experiment. Several clusters of genes with similar gene expression changes were observed (indicated as a1-2, b1-3, and c). (B) A heat map of a subset of genes from panel A, showing IFN-regulated genes and genes involved in inflammation. (C) A heat map of a one-way hierarchical cluster analysis of MVA transcriptome, using log2 RPKM, illustrating the temporal pattern of viral gene expression changes. (D) Gene set enrichment analyses (GSEA) showing differences of gene expression in several pathways including IFN-*γ*, IFN-α, inflammatory responses, and IL-6_JAK_STAT3 signaling. (E) Representative examples of heat-iMVA or live MVA-induced *Ifna4*, *Ifnb*, *Cd40*, and *Irf7* transcripts in WT and STING^Gt/Gt^ BMDCs.

MVA infection of BMDCs resulted in the temporal expression of viral RNAs, as shown by the unbiased hierarchical cluster analysis (figure 5C), consistent with published results of RNAseq of WT VAC-infected HeLa cells^37^. By contrast, heat-iMVA infection of cDCs did not result in significant levels of viral transcripts detected by the RNA-seq method (data not shown). Gene Set Enrichment Analyses (GSEA) confirmed that heat-iMVA-induced IFN-α, IFN-*γ*, inflammatory responses, and IL-6/JAK/STAT3 signaling in WT BMDCs but not in STING-deficient DCs (figure 5D). STING-dependent induction of *Ifna4, Ifnb1, Cd40, and Irf7* in heat-iMVA in BMDCs are shown in figure 5E. Together, RNA-seq analyses showed that heat-iMVA is more immune-stimulatory likely due to the lack of expression of viral inhibitory genes, and heat-iMVA-induction of type I IFN and IFN stimulated genes (ISGs) is largely dependent on STING.

### Heat-iMVA promotes migratory DC trafficking and maturation of resident DCs in the skin dLNs

The DC lineage is heterogeneous and composed of migratory and resident DCs ^38^. Migratory DCs capture antigens in the peripheral tissue and then mature, followed by migration to the draining lymph nodes, where they present antigens to naïve T cells. They can also transfer antigens to resident DCs ^39 40^. We analyzed various DC populations in skin dLNs after vaccination using a similar gating strategy as reported ^41^. First, we were able to confirm six distinct DC populations in the skin dLNs: (i) MHC-ΙΙ^+^CD11c^+^ migratory DCs and MHC-ΙΙ^Int^CD11c^+^ resident DCs; (ii) migratory DCs further separated into CD11b^+^ DCs, Langerin^-^ CD11b^-^ DCs, and Langerin^+^ DCs (CD103^+^ DCs and Langerhans cells); and (iii) resident DCs composed of CD8α^+^ lymphoid-resident DCs and CD8α^-^ lymphoid-resident DCs (Supplemental figure 3). Second, we tested which DCs subsets efficiently phagocytosing OVA antigen labeled with an Alexa Fluor 647 dye (OVA-647) and have the capacity to migrate to the skin dLNs. We intradermally injected OVA-647 into the right flanks of mice and harvested the skin dLNs at 24 h post-injection. Consistent with a previous report showing that migratory DCs are responsible for transferring antigens to skin draining LN ^42^, we observed that OVA-647 was mostly found in the three types of migratory DCs, including CD11b^+^, CD103^+^, and CD11b^-^CD103^-^ DCs, but rarely detected in resident DCs (figure 6A and 6C). Co-administration of OVA-647 with heat-iMVA increased the percentage of OVA-647^+^ CD11b^+^, CD103^+^, CD11b^-^CD103^-^, and CD8α^+^ DCs, compared with injection of OVA-647 alone (figure 6C and 6E). These results suggest that co-administration of OVA-647 with heat-iMVA enhances the capacity of migratory DCs to transport phagocytosed antigen to the skin dLNs and facilitates the antigen transfer from migratory DCs to CD8α^+^ DCs, a lymphoid-resident DC population critical for antigen cross-presentation. Migratory DCs expressed higher levels of CD86 maturation marker than resident DCs (figure 6B and 6D). Intradermal vaccination with OVA plus heat-iMVA induced higher levels of CD86 on resident DCs (CD8^+^ DC or CD8^-^ DC) than with OVA alone (figure 6D and 6F). By contrast, heat-iMVA co-administration did not change the maturation status of migratory DCs entering skin draining LNs (figure 6D and 6F). Our results indicate that intradermal co-administration of heat-iMVA with OVA antigen promotes antigen-carrying migratory DCs trafficking into the skin draining LN and transferring of antigens from migratory DCs to CD8α^+^ DCs and induce resident DC maturation.

**Figure 6.**
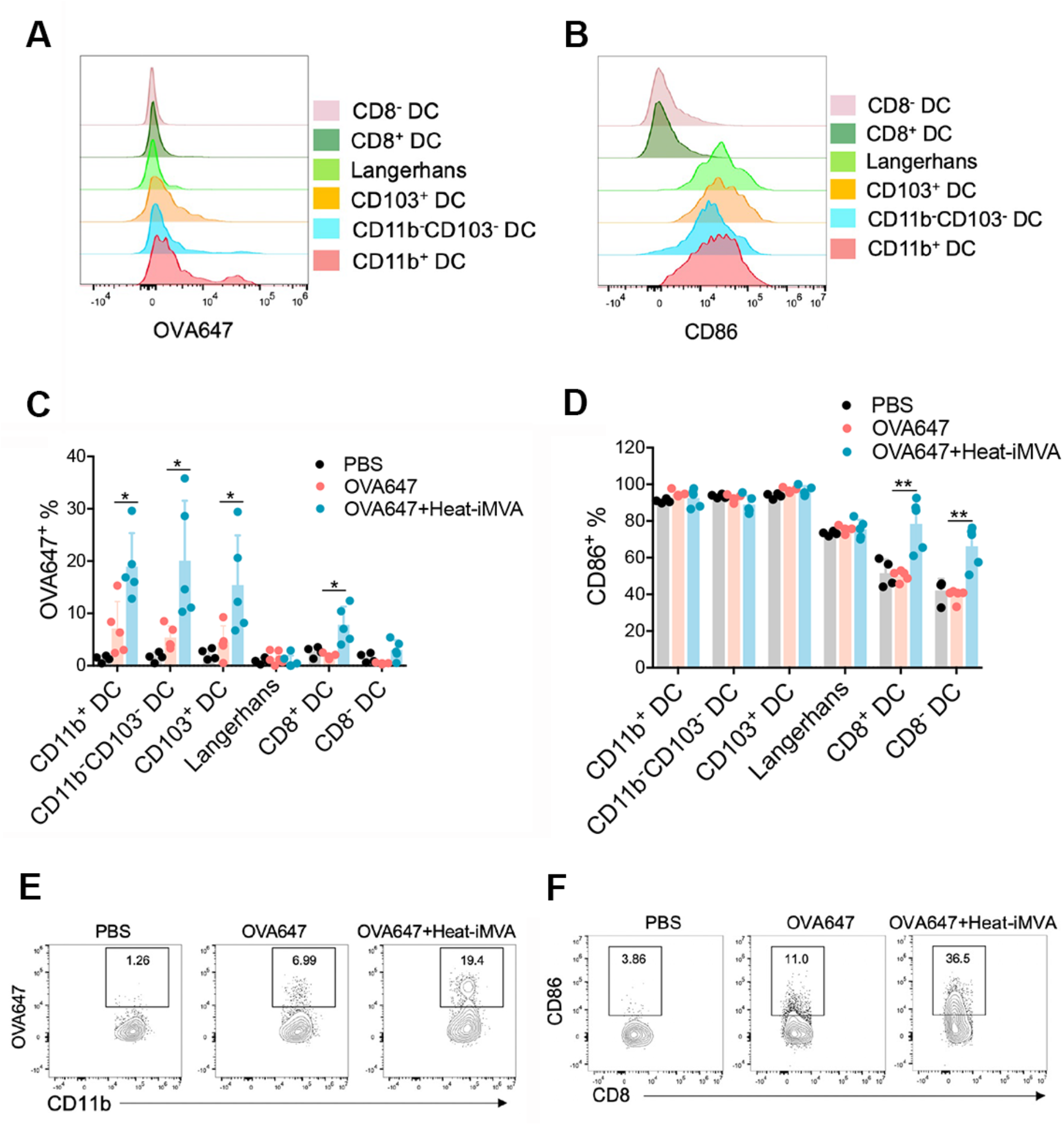
Co-administration of heat-iMVA promotes trafficking of antigen-carrying migratory DC into skin draining LN and activation of resident dendritic cells. C57/B6J mice were intradermally vaccinated with OVA647 (5 μg) in the presence or absence of heat-iMVA (10^7^ pfu). (A-B) After 24 h, OVA647 intensities in different dendritic cells populations from dLNs were measured. (C) Cell numbers of different DC populations were calculated. (D-E) After 24 h, CD86 expressions in different dendritic cells populations from dLNs were measured. (E-F) Representative dot plots of CD86 expression in CD8^-^ dendritic cells (E) and CD8^+^ dendritic cells (F). Data are represented as mean ± SEM (*n* = 3-5; \**P* < 0.05, and \*\**P* < 0.01; unpaired *t* test). Data are representative of three independent experiments.

### Co-administration of tumor neoantigen peptides with heat-iMVA improves antitumor effects in a murine therapeutic vaccination model

Here, we tested whether therapeutic vaccination with neoantigen peptides plus heat-iMVA would delay tumor growth in a murine B16-F10 melanoma model. Three days after B16-F10 cells were implanted, we subcutaneously co-administered melanoma neoantigen peptides (M27, M30, and M48) plus heat-iMVA twice, four days apart, and monitored tumor growth and mouse survival (figure 7A). Neoantigen peptides alone only minimally delayed tumor growth (figure 7B-7D).

**Figure 7.**
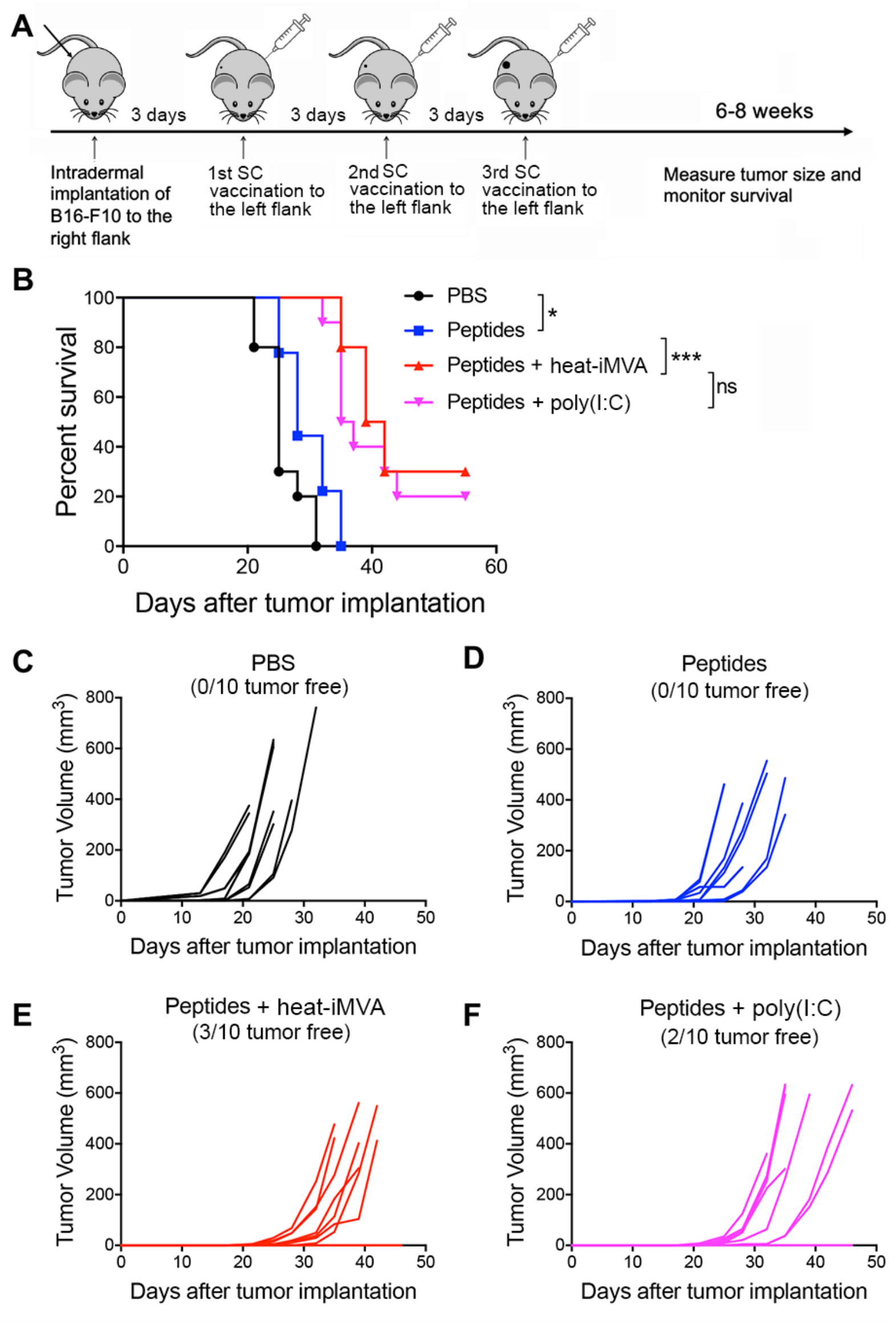
Combination of B16-F10 neoantigen peptides with heat-iMVA vaccination significantly increases the overall response and cure rates in a unilateral B16-F10 implantation model. (A) Tumor implantation and neoantigen peptide vaccination scheme in a unilateral B16-F10 tumor implantation model. 5 x 10^4^ B16-F10 were intradermally implanted into the right flanks of C57BL/6J mice. On day 3, 6, and 9, mice were vaccinated subcutaneously on the left flanks with B16-F10 neoantigen peptide mix (M27, M30, and M48) with or without the indicated adjuvants. (B) Kaplan-Meier survival curve of tumor-bearing mice treated with PBS, peptides (M27, M30, and M48, 100 μg/each), peptides plus heat-iMVA (an equivalent of 10^7^ pfu), or peptides plus poly(I:C) (50 μg) (*n* = 10, \**P* <0.05 and \*\*\**P* <0.001; Mantel-Cox test). (C, D, E, F) Tumor volumes over days after implantation in mice vaccinated with PBS (C), peptides (D), peptides + heat-iMVA (E), peptides + poly(I:C) (F). Data are representative of two independent experiments.

However, co-administration of neoantigen peptides with heat-iMVA cured B16-F10 melanoma in 30% of treated mice and prolonged the median survival (figure 7B, 7D, and 7E). Likewise, co-administration of poly(I:C) (50 μg) with neoantigen peptides also improved therapeutic vaccine efficacy with potency similar to heat-iMVA (figure 7B, 7D, and 7F). We did, however, observe side effects, including weight loss in the poly(I:C) group (but not with heat-iMVA). These results indicate that heat-iMVA could be a safe and potent vaccine adjuvant that eradicates or delays tumor growth in a murine therapeutic vaccination model.

### Heat-iMVA is a potent immune adjuvant for irradiated whole-cell vaccines

Use of irradiated whole-cell vaccines bypass the need to identify tumor-associated antigens or neoantigens and allows presentation of multiple tumor antigens for recognition by the host immune system. Here, mice were intradermally implanted with B16-F10 melanoma cells. After three days, mice were vaccinated subcutaneously with either irradiated B16-GM-CSF cells alone, irradiated B16-GM-CSF + heat-iMVA, or irradiated B16-GM-CSF + poly(I:C) on the contralateral flanks twice, four days apart (Supplemental figure 4A). Vaccination with irradiated B16-GM-CSF + heat-iMVA cured 30% of tumor-bearing mice and extended the median survival, which was more efficacious than using poly(I:C) as a vaccine adjuvant (Supplemental figure 4B and 4C). These results indicate that heat-iMVA is a potent vaccine adjuvant for irradiated whole-cell vaccination.

### Co-administration of heat-iMVA with SARS-CoV2 spike protein promotes robust neutralizing antibody production

Here, we tested whether co-administration of recombinant spike protein with heat-iMVA generates anti-spike neutralizing antibodies. Our results showed that vaccination with spike protein alone slightly induced anti-spike IgG1 and IgG2c antibodies (Supplemental figure 5A and 5B), while co-administration of spike protein + heat-iMVA increased IgG1 levels by 20-fold and IgG2c levels by 250-fold compared with vaccination with spike protein alone (Supplemental figure 5A and 5B). To investigate whether vaccination-induced antibodies could block SARS-CoV-2 infection, we performed a neutralization assay using a SARS-CoV-2 pseudovirus.

Without any pretreatment, the SARS-CoV-2 pseudovirus carrying the gene encoding GFP efficiently infected human ACE2-expressing HEK293T cells, as shown by the GFP^+^ cells (Supplemental figure 5C). Pretreatment with serum from the spike + heat-iMVA group at 1:100 dilution efficiently blocked SARS-CoV-2 pseudovirus infection, whereas pretreatment with serum from the spike alone group only weakly reduced pseudovirus infection (Supplemental figure 5C). Flow cytometry analysis of GFP^+^ cells confirmed our observation (Supplemental figure 5D). Serum neutralizing antibody titers in the two vaccination groups and a PBS-mock vaccination group was determined (Supplemental figure 5D and 5E). ID50 (50% inhibitory dose) was defined as the reciprocal of the serum dilution that caused a 50% reduction of GFP^+^ cells compared with mock-treated samples. The serum neutralizing antibody titers (ID50) from spike + heat-iMVA group were 10-fold higher than those from the spike alone group (Supplemental figure 5F). Overall, our results indicate that heat-iMVA boosts the production of neutralizing antibodies when combined with the recombinant spike protein from SARS-CoV-2.

## DISCUSSION

In this study, we explored the use of heat-iMVA as a vaccine adjuvant for protein- or peptide-based subunit vaccines against cancers and infectious agents. MVA is an approved vaccine against smallpox, and a potential vaccine vector with an excellent safety profile. However, MVA expresses many immune-suppressive genes. Heat-iMVA preserves the ability to enter DCs but fails to express viral genes, thus inducing much higher levels of type I IFN and ISGs than live MVA. Here, we demonstrated that co-administration of heat-iMVA with soluble proteins or peptides generates Th1-biased cellular and humoral immune responses superior to known adjuvants, including CFA and AddaVax. In a murine therapeutic vaccination model, co-delivery of heat-iMVA with three B16-F10 neoantigen peptides delayed tumor growth and cured 30% of tumor-bearing mice, with similar efficacy as poly(I:C), but with less toxicity. Taken together, our results support the use of heat-iMVA as a vaccine adjuvant.

DCs are essential for priming naïve T cells to generate adaptive immune responses, and therefore are the primary targets of vaccine adjuvants ^11 43–46^. RNA-seq analyses of host transcriptomes of DCs infected with either live MVA or heat-iMVA revealed heat-iMVA as a more potent STING agonist than live MVA, inducing large subsets of genes involved in type I and type II IFN and inflammatory responses. We previously showed that heat-iMVA infection of BMDCs induced DC maturation in a STING-dependent manner ^25^. Consistent with this, we now find that heat-iMVA infection of BMDCs promotes antigen cross-presentation, which requires STING. Barnowski et al. showed that the STING pathway contributes to the generation of vaccinia immunodominant B8-specific CD8^+^ T cell, but not to anti-OVA CD8^+^ T cell responses, after intraperitoneal vaccination with MVA expressing OVA ^47^. Together, these results show that the STING pathway is involved in poxvirus-induced antiviral adaptive immunity as well as the poxvirus-mediated adjuvant effect.

Using Batf3^-/-^ mice, we also demonstrated that heat-iMVA-boosted antigen-specific CD8^+^ T cell responses are dependent on cDC1s, also known as CD103^+^/CD8α^+^ DCs. However, heat-iMVA-boosted antigen-specific CD4^+^ T cell responses were not lost in Batf3^-/-^ mice, suggesting that cDC2s, also known as CD11b^+^ DCs, might be responsible for antigen presentation via MHC-II. KastenmÜller et al. investigated the role of Batf3-dependent CD103^+^/CD8α DCs in protein-TLR7/8 agonist conjugate and they found that CD8^+^ T cell responses were significant reduced in Batf3 KO mice, whereas CD4^+^ T cell responses were not affected ^48^. Interestingly, heat-iMVA-boosted antigen-specific antibody responses were not affected in Batf3^-/-^ mice. This finding is consistent with a recent report that migratory CD11b^+^ DCs (cDC2s) are required for priming T follicular helper (Tfh) cells, a subset of CD4^+^ T cells, for antigen-specific antibody production ^49^.

An increasing body of evidence indicates that STING agonists can function as potent vaccine adjuvants ^9 10 36 50–52^. To probe the role of the STING-mediated cytosolic DNA-sensing pathway in heat-iMVA adjuvanticity, we used STING^Gt/Gt^ mice, which lack functional STING^53^. Our results demonstrate that STING contributes to the generation of antigen-specific IFN-*γ*^+^ CD8^+^ and CD4^+^ T cells and IgG2c antibody production potentiated by heat-iMVA. Given the essential roles of cDC1 and cDC2 in mediating CD8^+^ and CD4^+^ T cell priming, we surmise that STING signaling in cDC1 and cDC2 is important for heat-iMVA-induced immunogenicity. We note three major differences between heat-iMVA and small-molecule chemical STING agonists: (i) heat-iMVA enters cells naturally ^24^, whereas chemical STING agonists need lipophilic mediators to facilitate their entry; (ii) heat-inactivated vaccinia can also trigger type I IFN production in pDCs via a TLR7/TLR9/MyD88-dependent pathway ^24 26^, whereas chemical STING agonists are specific to the STING pathway; and (iii) heat-iMVA activates other danger signals, including the Absent In Melanoma 2 (AIM2) inflammasome (^54^ and data not shown), which might also be important for heat-iMVA adjuvanticity. Our results showed that SC heat-iMVA co-administration with OVA-647 significantly increased OVA-647^+^ migratory DCs in the dLNs. Interestingly, heat-iMVA also increased OVA-647^+^ CD8α^+^ DCs in the dLNs. Although migratory DCs in the dLNs exhibited high CD86 expression with or without heat-iMVA as a vaccine adjuvant, heat-iMVA co-delivery resulted in higher expression of CD86 on resident DCs, including both CD8^+^ DCs and CD8^-^ DCs. These results suggest that heat-iMVA co-administration promotes peripheral DC maturation and migration into dLNs, as well as LN-resident DC maturation. We speculate that some heat-iMVA virions (together with OVA-647^+^) might be transported, via the LN conduits, to the LN interior to resident DCs, as demonstrated for subcutaneously injected vaccinia virus or MVA ^55^.

Limitations of the study include but are limited to the following: (i) vaccination studies were performed in female C57BL/6J mice at 6-8 weeks of age. Potential age, sex, and species bias cannot be excluded; Because the Batf3 and STING-deficient mice are in C57BJ/6J background, we chose this strain for our study; (ii) we did not analyze what cell populations are infected by heat-iMVA after vaccination, because heat-iMVA does not produce viral proteins; we plan to do more sophisticated analyses with immune-activating recombinant MVA expressing fluorescent markers in our future studies; and (iii) we did not explore whether antigen conjugation to heat-iMVA or mixing antigen with Addavax plus heat-iMVA would further enhance immune efficacy in this study.

Nörder et al. investigated whether MVA could be used as a vaccine adjuvant, and they found that IM co-administration of MVA and OVA enhances the generation of antigen-specific antibody and T cell responses ^56^. MVA is non-replicative in most mammalian cells and heat-inactivation makes it safer and more immunogenic.

In conclusion, we envision that heat-iMVA can be used as a vaccine adjuvant for neoantigen-based or irradiated whole cell-based cancer vaccines based on its safety and immunogenicity. Heat-iMVA activation of STING signaling contributes to its adjuvanticity and heat-iMVA promotes antigen cross-presentation by Batf3-dependent CD103^+^/CD8α^+^ DCs to induce antigen-specific CD8^+^ T cell responses. Future work will focus on identifying viral inhibitors of the cGAS/STING pathway encoded by the MVA genome and engineering recombinant MVA to improve its immunogenicity and adjuvanticity.

## Acknowledgements

We thank the Flow Cytometry Core Facility and Molecular Cytology Core Facility at the Sloan Kettering Institute. We also thank the Rockefeller University Genomics Resource Center. We thank Peihong Dai’s technical support for BMDC RNA-seq experiments. We also thank Shuaitong Liu and Yi Wang’s technical assistance on antigen cross-presentation assay.

## Availability of data and materials

All data published in this report are available on reasonable request.

RNA-sequencing data have been deposited at NCBI Short-Read Archive (SRA) and are publicly available as of the date of publication under the BioProject number PRJNA743347.

https://dataview.ncbi.nlm.nih.gov/object/PRJNA743347?reviewer=23jdtbg40kcljcqf9b75ubqei6

## Authors’ contributions

N.Y. and L.D. designed and performed the experiments, analyzed the data, and prepared the manuscript. A.G. and C.M. performed the library preparation for RNA-seq of virus-infected BMDCs and analyzed the RNA-seq data. Memorial Sloan Kettering Cancer Center filed a patent application for the Heat-inactivated vaccinia virus as a vaccine immune adjuvant. J.D.W., T.M, and T.T. assisted in experimental design and data interpretation. All authors are involved in manuscript preparation. L.D. provided overall supervision of the study.

## Patient consent for publication

N/A

## Ethics approval and consent to participate

N/A

