## Supplemental files for "Heat-inactivated modified vaccinia virus Ankara boosts Th1-biased cellular and humoral immune responses as a vaccine adjuvant by activating the STING-mediated cytosolic DNA-sensing pathway"

Short title: heat-inactivated MVA as a vaccine adjuvant

This file contains:

- Supplemental figure 1
- Supplemental figure 2
- Supplemental figure 3
- Supplemental figure 4
- Supplemental figure 5

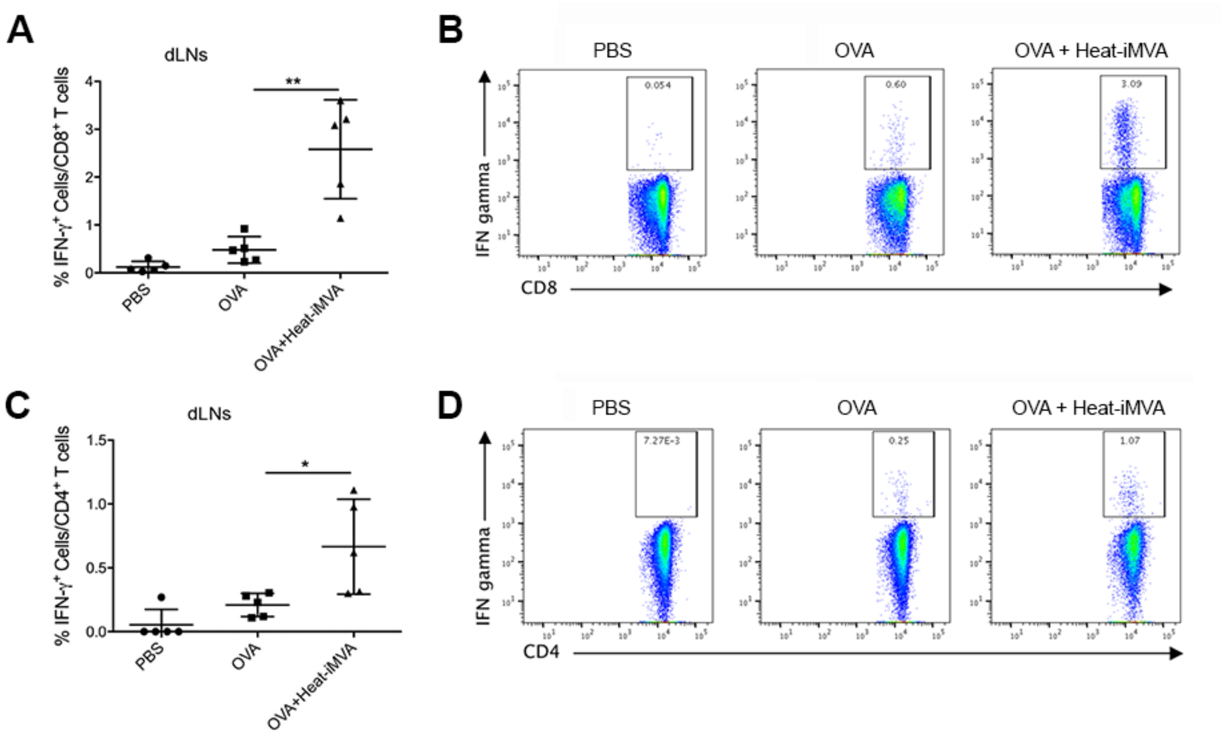


**Supplemental figure 1. Heat-inactivated MVA (Heat-iMVA) enhances antigen-specific T cell in the draining lymph nodes (dLNs) after intramuscular (IM) vaccination with chicken ovalbumin (OVA).** WT C57BL/6J mice were vaccinated on day 0 and day 14 with OVA (10 µg) or OVA (10 µg) plus Heat-iMVA ($10^7$ pfu) intramuscularly. On day 21, cells from dLNs from the vaccinated mice (A, B, C, D) were stimulated with $OVA_{257\text{-}264}$ (A, B) or $OVA_{323\text{-}339}$ peptide (C, D). The expressions of IFN-γ by $CD8^+$ T cells or $CD4^+$ T was measured by flow cytometry. (A) A representative graph of percentages of IFN-$\gamma^+$$CD8^+$ T cells in the dLNs of PBS, OVA, or OVA + Heat-iMVA-vaccinated mice. (B) Dot plots of IFN-$\gamma^+$$CD8^+$ T-cells in the dLNs. (C) A representative graph of percentages of IFN-$\gamma^+$$CD4^+$ T cells in the dLNs of PBS, OVA, or OVA + Heat-iMVA-vaccinated mice. (D) Dot plots of IFN-$\gamma^+$$CD4^+$ T-cells in the dLNs. Data are represented as mean ± SEM ($n$ = 3-5; $^*P < 0.05$, $^{**}P < 0.01$ and $^{***}P < 0.001$; unpaired $t$ test). Data are representative of three independent experiments.

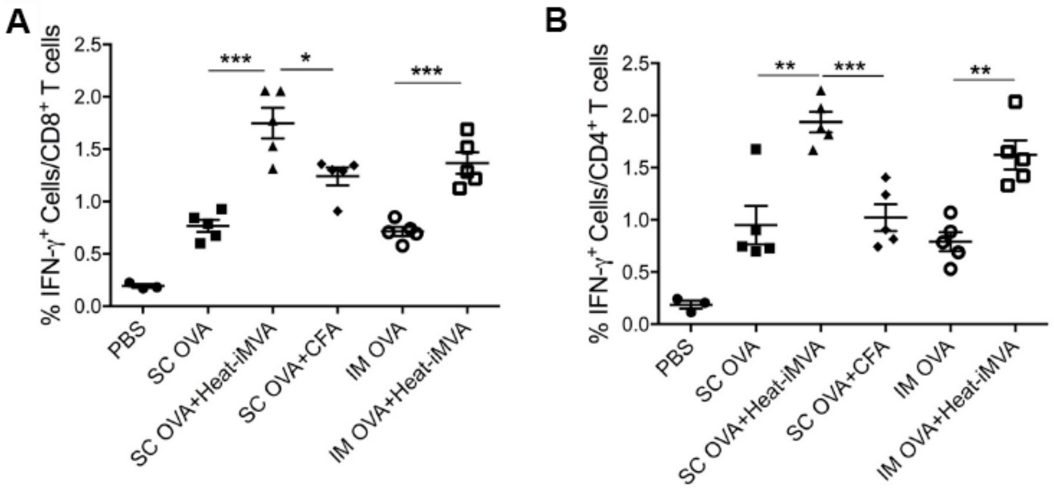


**Supplemental figure 2. Heat-iMVA induces stronger antigen-specific $CD8^+$ and $CD4^+$ T cell responses in the draining lymph nodes than complete Freund adjuvant (CFA) after subcutaneous vaccination.** Antigen-specific T cell responses were measured one week after intramuscular (IM) or subcutaneous (SC) vaccination on day 0 and day 14 with OVA (10 μg) in the presence or absence of Heat-iMVA (an equivalent amount of $10^7$ pfu) in C57BL/6J mice. CFA was also used as an adjuvant in SC vaccination. (A) On day 21, cells from the dLNs were stimulated with $OVA_{257-264}$ (SIINFEKL) peptide (10 μg/ml) for 12 h. The expression of IFN-γ by $CD8^+$ T-cells was measured by flow cytometry. (B) Cells from the dLNs were stimulated with $OVA_{323-339}$ (ISQAVHAAHAEINEAGR) peptide (10 μg/ml) for 12 h. The expression of IFN-γ by $CD4^+$ T-cells was measured by flow cytometry. Data are represented as mean ± SEM ($n$ = 3-5; $^{*}P < 0.05$, $^{**}P < 0.01$ and $^{***}P < 0.001$; unpaired $t$ test). Data are representative of two independent experiments.

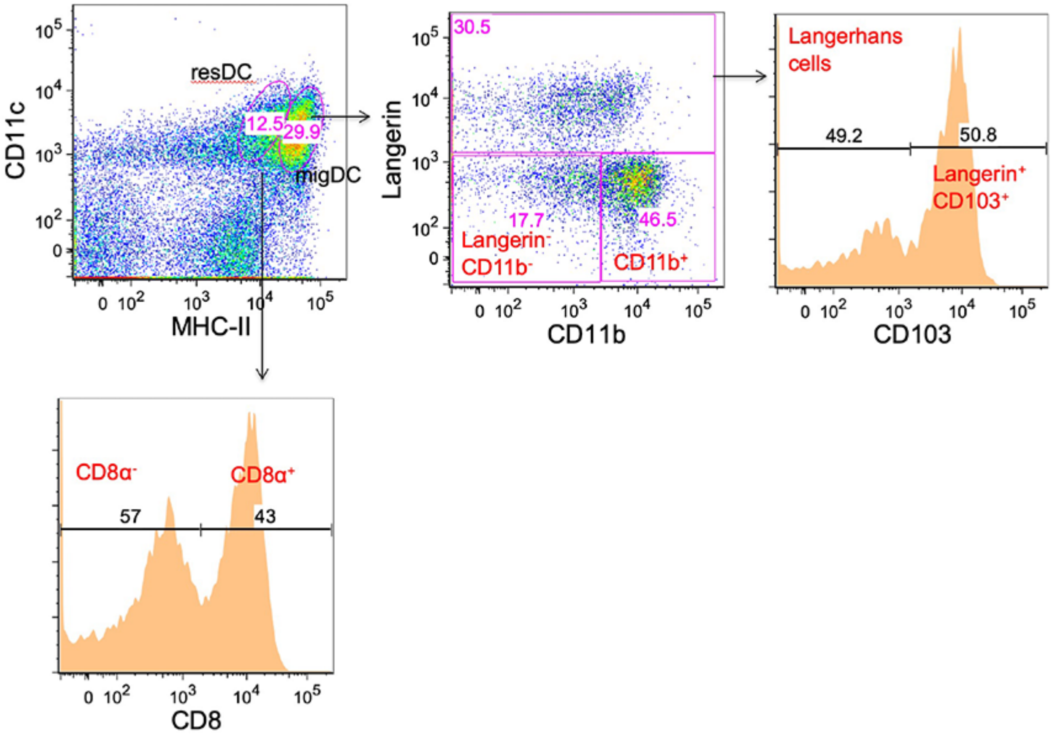


**Supplemental figure 3. Gating strategy of dendritic cell populations in skin LN.** Skin LN were harvested and digested into single-cell suspension. Cells were stained with a cocktail of antibodies to distinguish DC subsets. Within single cells, dead cells, CD19$^+$, DX5$^+$, TER119$^+$, and CD3$\varepsilon^+$ cells were excluded for analysis. The CD11c$^{hi}$ MHC II$^{int}$ population represents lymphoid resident DCs and can be further divided into CD8$^+$ and CD8$^-$ DCs. CD11c$^{int/hi}$ MHC II$^{hi}$ cells were further analyzed for the expression of Langerin and CD11b to define skin migratory DC subsets. Langerin$^-$ cells were divided into CD11b$^+$ DCs and CD11b$^-$ DCs. Langerin$^+$ cells were further analyzed for the expression of CD103 and divided into two groups: CD103$^+$ DCs and Langerhans cells.

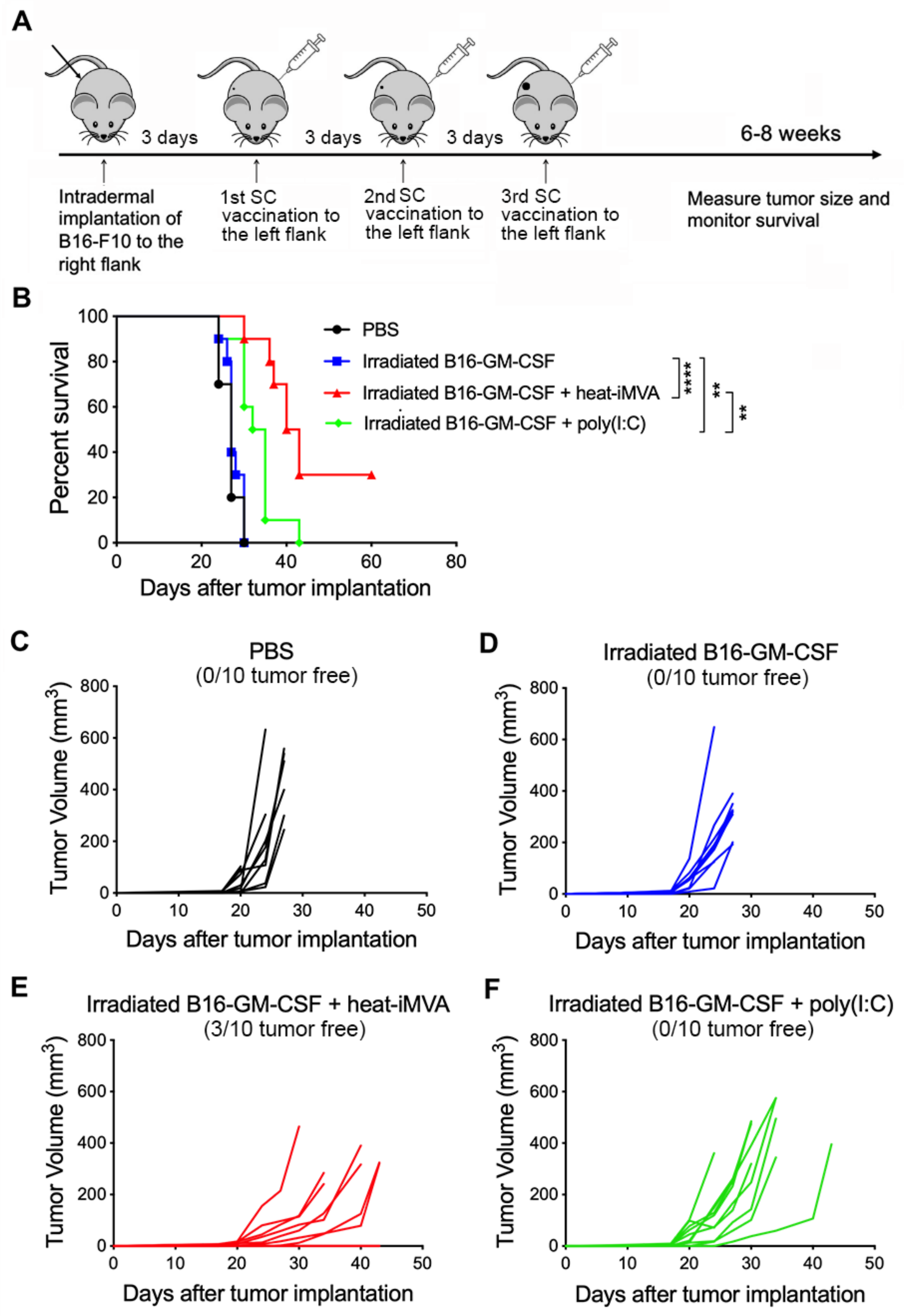
A
3 days
3 days
3 days
6-8 weeks
Intradermal implantation of B16-F10 to the right flank
1st SC vaccination to the left flank
2nd SC vaccination to the left flank
3rd SC vaccination to the left flank
Measure tumor size and monitor survival
B
Percent survival
Days after tumor implantation
PBS
Irradiated B16-GM-CSF
Irradiated B16-GM-CSF + heat-iMVA
Irradiated B16-GM-CSF + poly(I:C)
****
**
**
C
PBS
(0/10 tumor free)
Tumor Volume (mm3)
Days after tumor implantation
D
Irradiated B16-GM-CSF
(0/10 tumor free)
Tumor Volume (mm3)
Days after tumor implantation
E
Irradiated B16-GM-CSF + heat-iMVA
(3/10 tumor free)
Tumor Volume (mm3)
Days after tumor implantation
F
Irradiated B16-GM-CSF + poly(I:C)
(0/10 tumor free)
Tumor Volume (mm3)
Days after tumor implantation

**Supplemental figure 4. Combination of irradiated B16-GM-CSF with Heat-iMVA vaccination is more effective in preventing or delaying tumor growth compared with poly(I:C) in a unilateral B16-F10 tumor implantation model.** (A) Tumor implantation and irradiated tumor whole-cell vaccination scheme in a unilateral B16-F10 tumor implantation model. 5 x $10^4$ B16-F10 were intradermally implanted into the right flanks of C57BL/6J mice. On day 3, 6, and 9, mice were vaccinated subcutaneously on the left flanks with 1 x $10^6$ irradiated B16-GM-CSF cells with or without the indicated adjuvants. (B) Kaplan-Meier survival curve of tumor-bearing mice treated with PBS, irradiated B16-GM-CSF ($10^6$ cells) alone, irradiated B16-GM-CSF ($10^6$ cells) + heat-iMVA (an equivalent of $10^7$ pfu) or irradiated B16-GM-CSF ($10^6$ cells) + poly(I:C) (5 μg) ($n = 10$, $^{**}P < 0.01$ and $^{****}P < 0.0001$; Mantel-Cox test). (C, D, E, F) Tumor volumes over days after implantation in mice vaccinated with PBS (C), irradiated B16-GM-CSF (D), irradiated B16-GM-CSF plus heat-iMVA (E), irradiated B16-GM-CSF plus poly(I:C) (F). Data are representative of two independent experiments.

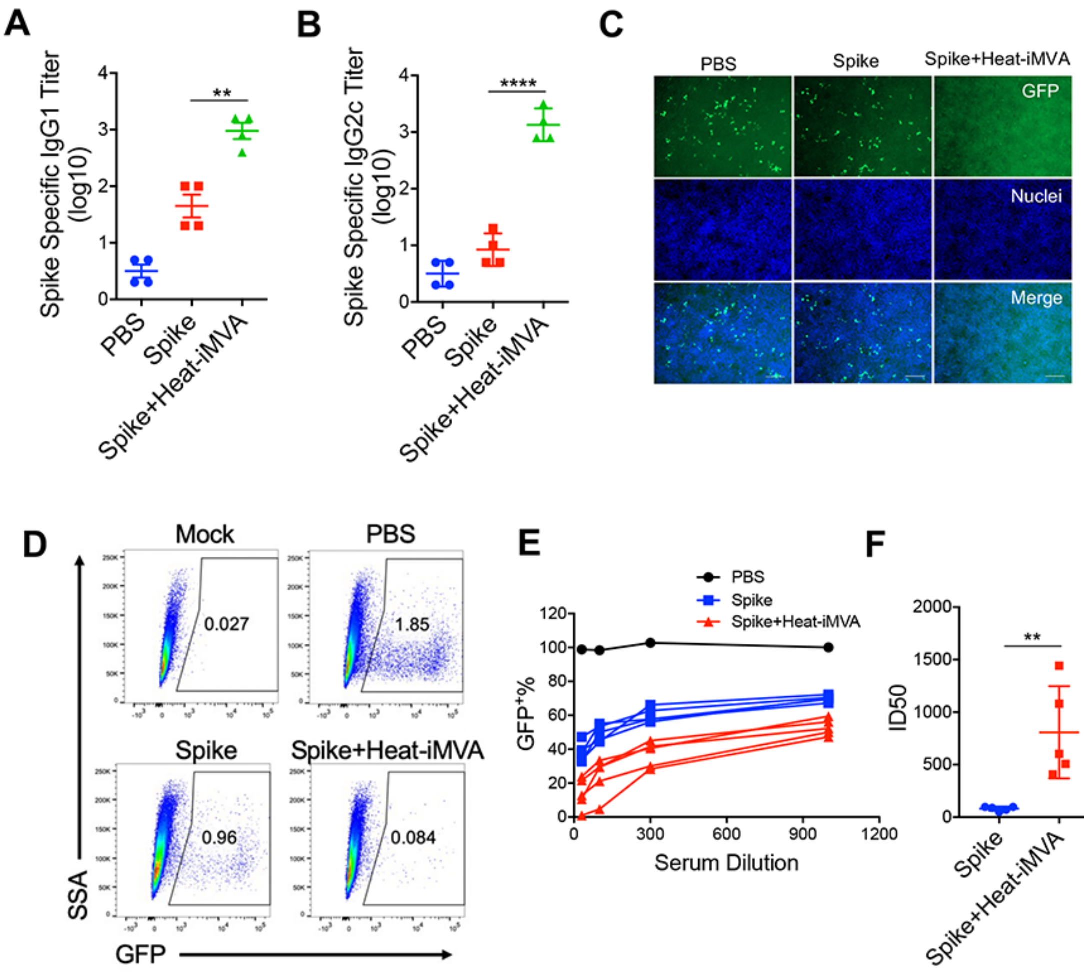


**Supplemental figure 5. Heat-iMVA promotes stronger Th1 responses and IgG1, IgG2c production after intramuscular (IM) vaccination with SARS-CoV2 full length spike protein.** WT C57BL/6J mice were vaccinated on day 0 and day 21 with SARS-CoV2 Spike (1 µg) or Spike (1 µg) plus Heat-iMVA ($10^7$ pfu) intramuscularly. (A-B) Spike-specific immunoglobulin G1 (IgG1) or immunoglobulin G2c (IgG2c) titers in the serum from PBS, Spike, or Spike plus Heat-iMVA-vaccinated mice one week after second vaccination were determined by ELISA. (C-F) HEK293T-ACE2 cells were infected with SARS-CoV-2 pseudovirus at the presence of mouse serum (1:100 dilution). After 48 h, spike protein mediated virus entry was detected by GFP expression. (C) GFP was observed by fluorescence microscope. (D) GFP was measured by FACS analysis. (E) The neutralizing antibodies in serum at different dilution was detected by GFP expression based on FACS analysis. (F) 50% inhibitory dose (ID50) was determined. Data are represented as mean ± SEM ($n$ = 4-5; $^{**}P < 0.01$ and $^{****}P < 0.0001$; unpaired $t$ test). Data are representative of two (A-B) or three (C-F) independent experiments.
